# A Graph Coarsening Algorithm for Compressing Representations of Single-Cell Data with Clinical or Experimental Attributes

**DOI:** 10.1101/2022.07.30.502142

**Authors:** Chi-Jane Chen, Emma Crawford, Natalie Stanley

## Abstract

Graph-based algorithms have become essential in the analysis of single-cell data for numerous tasks, such as automated cell-phenotyping and identifying cellular correlates of experimental perturbations or disease states. In large multi-patient, multi-sample single-cell datasets, the analysis of cell-cell similarity graphs representations of these data becomes computationally prohibitive. Here, we introduce *cytocoarsening*, a novel graph-coarsening algorithm that significantly reduces the size of single-cell graph representations, which can then used as input to downstream bioinformatics algorithms for improved computational efficiency. Uniquely, cytocoarsening considers both phenotypical similarity of cells and similarity of cells’ associated clinical or experimental attributes in order to more readily identify condition-specific cell populations. The resulting coarse graph representations were evaluated based on both their structural correctness and the capacity of downstream algorithms to uncover the same biological conclusions as if the full graph had been used. Cytocoarsening is provided as open source code at https://github.com/ChenCookie/cytocoarsening.

## 1. Introduction

Advancements in a range of single-cell technologies, such as flow and mass cytometry and single-cell RNA sequencing, have become essential in uncovering and understanding cellular heterogeneity in a range of translational applications.^1–3^ These immune profiling techniques have proven to be particularly essential in unraveling immunological heterogeneity through the simultaneous measurement of 20-45 protein markers in each cell.^4^ This simultaneous measurement enables both phenotypic (e.g. cellular identity) and functional characterization of cells.^5^ Despite effective identification and characterization of immune cell-types, a current challenge is to accurately link these immune cells to external *attributes* of interest, such as clinical labels or experimental perturbations.^6–9^ For example, it is common in translational applications to profile blood samples from patients *across* clinical phenotypes or disease states in order to identify the driving, stratifying cell-types.^6,10^ Blood samples are also often perturbed through stimulation,^11^ and cellular correlates are identified by observing functional responses to the stimulation. Moreover, to efficiently link cellular heterogeneity to clinical or experimental attributes, automated bioinformatics methods have become critical in analysis.

Many of the bioinformatics algorithms for such tasks operate on a graph representation of the single-cell data.^7–9^ In these graphs, nodes are cells, and edges between a pair of cells imply that they are sufficiently similar across measured features (for example, the aforementioned protein markers). The task at hand is to use the graph structure to identify cells that are prototypical of particular external attributes, such as clinical or experimental labels. MELD^7^ accomplishes this by modeling the external attributes as a signal on the graph and computing a score for each cell reflecting its probability of association with each condition. To exemplify another approach, Milo^8^ and CNA^9^ seek to identify critical *cellular neighborhoods*, or groups of phenotypically-similar cells enriched across attributes.

Practically, it is challenging to apply these bioinformatics algorithms to the extremely large graph representations of multi-patient, multi-sample cohorts with millions of cells. Although the large graph size would make computations on it prohibitive, the graph inherently involves redundant information, since we have multiple cellular instances from a single population encoding the same biological information. To reduce the graph size, then, we merge redundant cells into *coarse nodes* or *super nodes*, leveraging existing graph-coarsening strategies^12,13^ and adapting them to consider biologically relevant external attributes. The rich literature of existing graph-coarsening methods^13–18^ tend to optimize for merges of nodes that maintain critical structural and spectral properties for the original graph, but do not consider these node attributes.

As an example of a graph-coarsening approach, Loukas *et al*. proposed a family of *local variation* algorithms to simplify and reduce the size of the original graph.^14^ These algorithms begin with a family of *coarsening candidate sets*: subsets of nodes that are known to be highly related based on the graph structure. The two main approaches discussed are edge-based variation (LV-E) or node-based variation (LV-N). Using LV-E, the candidate sets are exactly the edge pairs of the graph. In contrast, the candidate sets in LV-N are formed by grouping each node with its immediate neighborhood. In Ref.14, Loukas *et al*. compared these variation-based methods to other graph coarsening methods, including heavy-edge matching (HEM),^15^ algebraic distance (AD),^16^ and affinity (AFF).^17^ The local variation methods outperformed these methods in spectral approximation, and all of the methods (with the exception of AFF, which is slower) scale quasi-linearly in the number of edges in the graph. Briefly, HEM seeks to coarsen the graph such that the principal eigenvalues and eigenspaces of the coarsened graph Laplacian are close to those of the original graph Laplacian. Instead of considering spectral properties, the AD and AFF methods identify nodes to merge by considering the connectedness of both individual nodes and node neighborhoods.

With existing coarsening approaches focusing primarily on preserving overall graph structure or underlying spectral properties, we seek to adapt the methods to additionally take into account external attributes of the cells, such as clinical state or experimental perturbation status. Our method will therefore merge individual nodes (representing cells) into coarse nodes according to both cellular phenotype and associated attributes (see overview figure, Fig 1). This gives us a graph of reduced size to use as input for downstream bioinformatics algorithms, and it facilitates simpler identification of cells that are related both in phenotype and in clinical or experimental attribute.

**Fig. 1.**
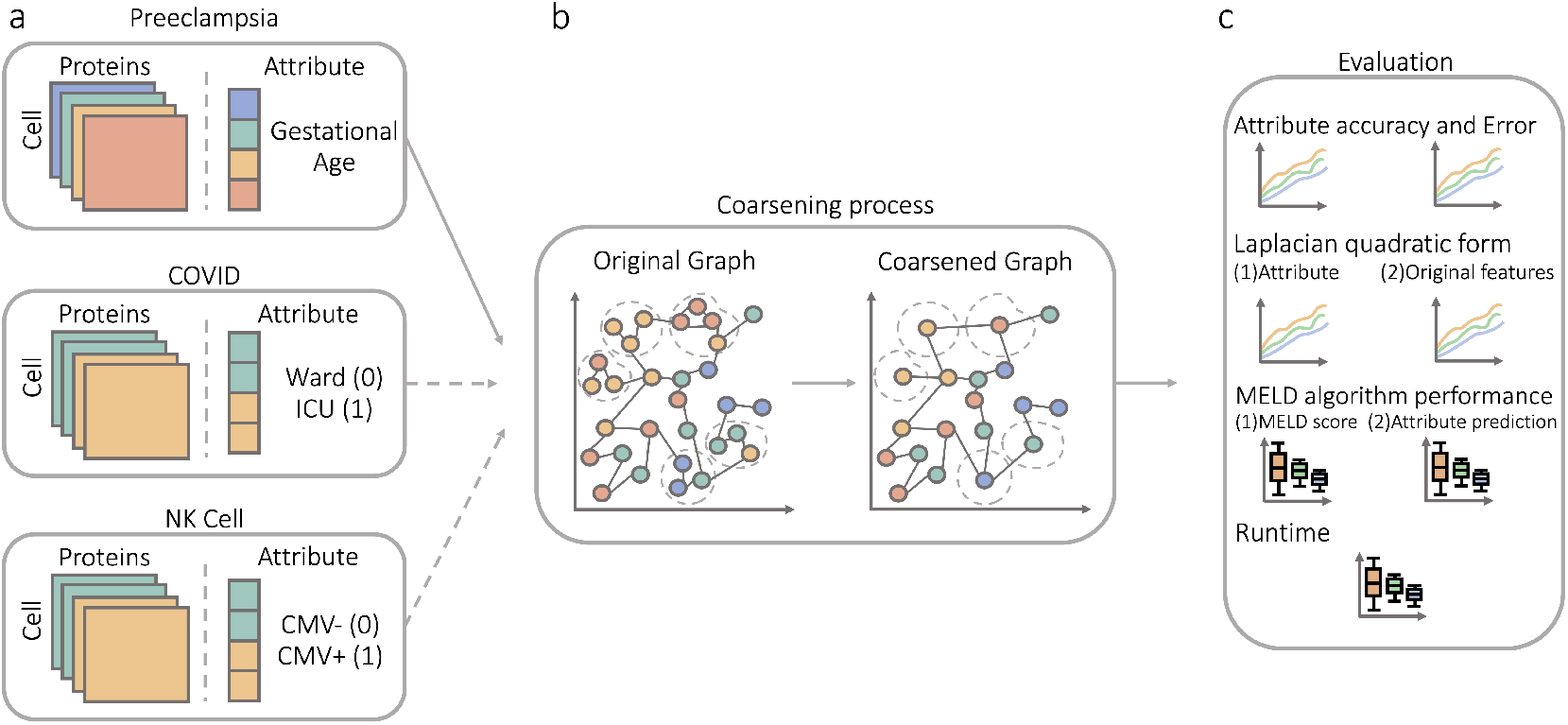
Overview. Given a multi-sample single-cell dataset with clinical attributes (**a**), the cytocoarsening algorithm creates a *coarse* graph representation of all cells (**b**). The coarse graph representation takes into account phenotypic similarity of cells (edges) and the clinical attributes (colors). (**c**) Quantitative evaluation metrics were developed to assess the quality of the coarse graph representation and its effectiveness as input to downstream graph-based bioinformatics algorithms.

## 2. Methods

### Notation and problem formulation

We consider a multi-sample single-cell dataset with *p* profiled samples, denoted as 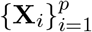. Here, each 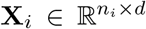 represents the *d* protein or gene expression measurements for each of the *n*_*i*_ cells measured in sample *i*. We also assume that each cell has an *attribute label* (such as experimental label or disease state), encoded in the vector **x**. A graph representation of all of these cells would render further computation expensive and time-consuming. Thus, we seek a graph representation of the 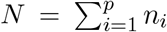 cells that has *N* ^′^ *<< N* nodes while still representing the biologically relevant information that would be present in the full graph. To accomplish this, we introduce the *cytocoarsening* algorithm. In this section, we outline the general steps of the algorithm; pseudocode is provided in Algorithm 1.

### Graph representation of single-cell data

The algorithm begins by constructing a joint graph representation 𝒢 of all profiled cells across samples. Given a data matrix of cells × measured features defined as **X** = [**X**_1_|**X**_2_| … |**X**_*p*_] (where | denotes vertical concatenation), each cell is connected to its *K* nearest neighbors in the measured feature space (`KNN()` in Algorithm 1). To actually carry out computations with this graph, we will use the adjacency matrix **A**, which has all the edge weights of the graph encoded in its off-diagonal entries and zeros on the diagonal. We will also use the graph Laplacian **L**, which is exactly the negative of this matrix but with a diagonal instead defined as 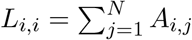.

### Establishing and ranking coarsening candidates

The KNN graph is used to define the *coarsening candidate node sets* as each node and its *K* nearest neighbors; the candidate sets are stored in the list *C* with corresponding index set list *I*^*C*^, i.e. 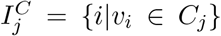 (KNN enumeration → `get.K.Neighborhoods`(), indices of nodes within coarsening candidate → `get.Index.Sets`() in Algorithm 1). To decide which candidate sets to coarsen, we define two different cost functions: distance in feature space (**c**^*d*^) and graph-level attribute variation (**c**^*q*^).

#### Algorithm 1

Cytocoarsening

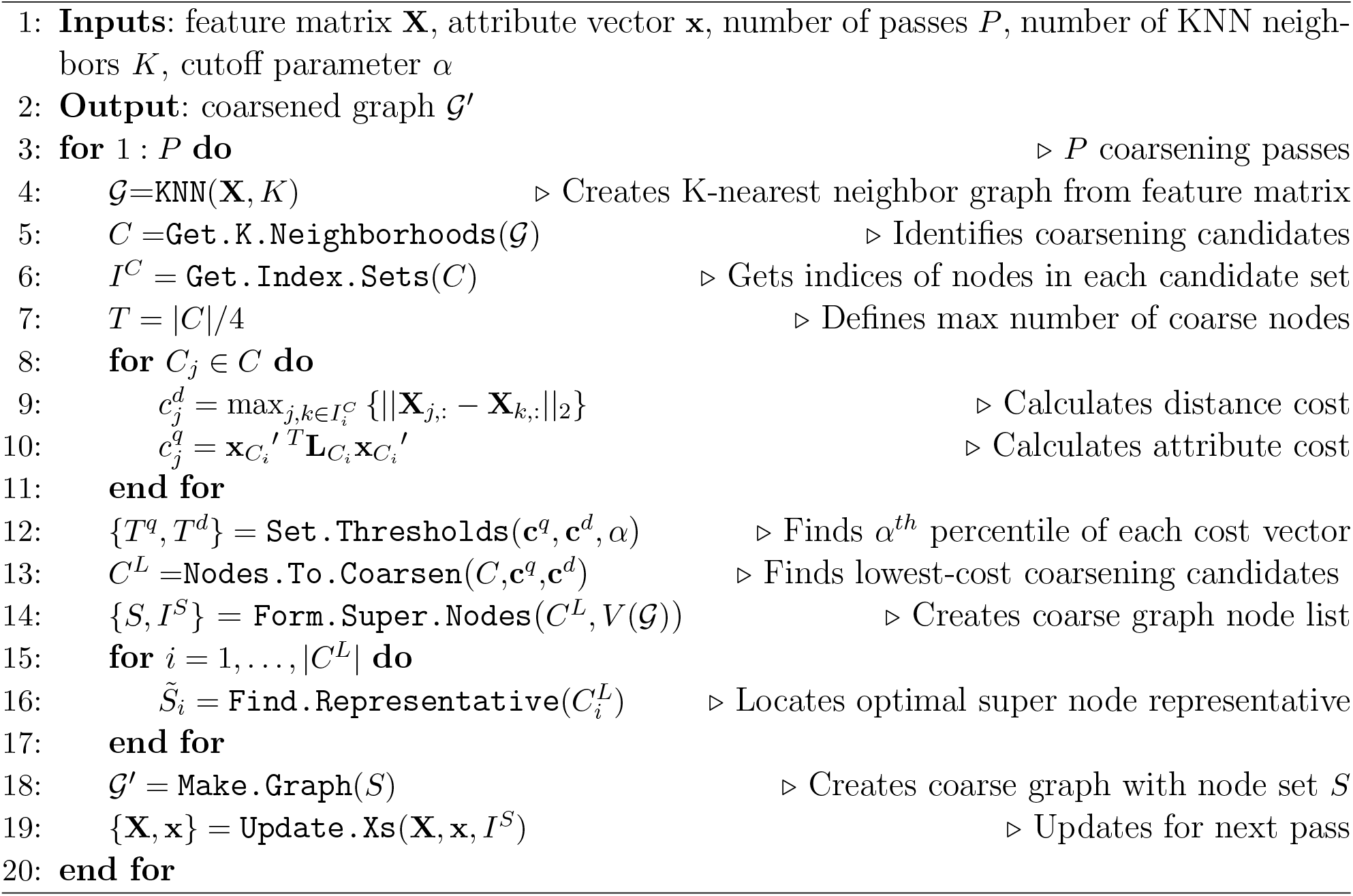

### Distance cost (c^*d*^)

The distance cost reflects the overall phenotypical similarity between cells in a coarsening candidate to ensure that highly similar nodes are likely to be aggregated. We define 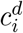, the distance cost of the *i*^*th*^ coarsening candidate, as the maximum euclidean distance of all cells within a coarsening candidate:

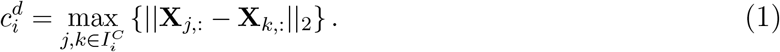

### Attribute cost (c^*q*^)

The attribute cost measures the overall variation of the attributes of cells within a coarsening candidate, so that we can prioritize merges of cells with similar attributes. Given a coarsening candidate, *C*_*i*_, we can extract its sub-adjacency matrix, 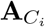 via 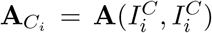 and compute its corresponding Laplacian matrix, 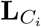. We further let 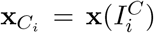 be the corresponding subvector of attributes for the coarsening candidate set. Then the attribute cost 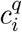 for coarsening candidate *C*_*i*_ is computed as

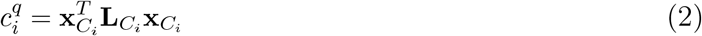

### Joint cost (r)

We use a joint ranking criteria to rank coarsening candidates according to their phenotypic between-cell similarity (**c**^*d*^) and attribute consistency (**c**^*q*^) by simply taking the log of their geometric mean:

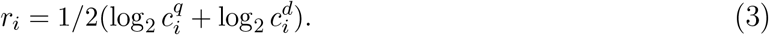

The 30 coarsening candidates with the lowest joint cost are then considered for further evaluation.

### Evaluating coarsening candidates

A coarsening candidate *C* will be added to the coarsening list *C*^*L*^ (i.e. selected to be aggregated) if all of the following are true: 1) less than *T* coarsening candidates have been chosen, 2) both costs **c**^*q*^ and **c**^*d*^ are below some percentile thresholds *T*^*q*^ and *T*^*d*^ (see `Set.Thresholds()` in Algorithm 1) to make sure both two costs are sufficiently low, 3) none of the nodes in *C* are already represented in the coarsening list. Our method will stop trying to find more coarsening candidates to merge if all coarsening candidates remaining have a cost larger than *c*_*max*_, a global constant. If some nodes in the candidate are already present in *C*^*L*^, then those nodes are removed from the set and the costs are recomputed for this smaller candidate set. In the cases where only one node remains or there are no edges between the remaining candidate nodes, we assign both costs the value of *c*_*max*_ in order to remove that set from consideration (see function `Nodes.To.Coarsen`() in Algorithm 1). Once the coarse node sets have been decided, we form the node set for the coarse graph *S* (with corresponding index set *I*^*S*^) by taking the union of the coarse nodes with all the individual nodes from the original graph (see function `Form.Super.Nodes`() in Algorithm 1).

### Defining super node representatives

Once we know which sets of nodes to merge, we find the original node in each set that is most representative of the group by considering two factors: phenotypical similarity and attribute similarity. Consider the *i*^*th*^ super node in the following discussion. For phenotypical similarity, we find the mean point of the nodes in feature space 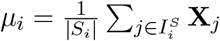, and then we calculate the euclidean distance from *µ*_*i*_ to each node in the set. Weights are assigned so that nodes closer to *µ*_*i*_ are more highly weighted. For attribute similarity, we sort the attribute labels by the number of their occurrences in *S*_*i*_ and weight the nodes so that nodes with frequently-occurring attribute values are more highly weighted. To combine these two weights, we normalize them individually and add them together. The representative node is then chosen as the one with the maximum aggregate weight. We will denote the representative node for the *i*^*th*^ super node as 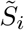, with original graph index 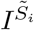 (see function `Find.Representative`() in Algorithm 1).

### Updating edge list

An edge is defined between a pair of nodes *S*_*i*_ and *S*_*j*_ in the coarse graph if, in the original graph, there was at least one edge between any of the nodes in *S*_*i*_ and *S*_*j*_. (`Make.Graph`() function in Algorithm 1).

The above outlines one pass of the algorithm. To coarsen further, we update the feature matrix 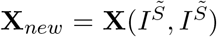 and the attribute vector 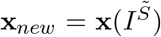 (see function `Update.Xs`() in Algorithm 1).

## 3. Results

To explore the effects of graph coarsening on biological information, we applied our cytocoarsening algorithm to three publicly available mass cytometry (e.g. CyTOF) datasets. First, the *preeclampsia dataset* ^19^ profiles blood samples collected 9.7 millions cells from 45 women throughout their pregnancies (33 features measured per cell). The clinical attribute of interest for this dataset was cell gestational age, which ranged from 8 to 28 weeks. Next, the *covid dataset* ^20^ contains 6.5 million cells collected from 49 total patients (23 features measured per cell). The patients ranged in severity with 6 healthy patients, 23 patients having mild cases of COVID, and 20 experiencing severe responses and were under ICU care. Due to the imbalance in the number of patients for each severity level, we only considered cells from 22 mild patients (one sample had less than 1,000 cells and was thus not considered) and 20 patients that had severe (ICU) COVID. The attribute of interest was disease severity (mild or severe). Finally, the *NK-cell dataset* ^21^ contains 261 thousand cells collected from 20 total patients (29 measured features per cell). Cytomegalovirus (CMV) status was the attribute of interest, with nine patients being positive for Cytomegalovirus (CMV) and 11 being negative for CMV.

We performed several experiments (Fig. 1c) on cytocoarsening and existing coarsening methods (LV-E, LV-N, HEM, AD, and AFF^14^) to quantify their effectiveness in preserving structural and attribute information and in acting as input to downstream graph-based bioinformatics tasks. All experiments were repeated 30 times, sampling a new subset of cells from each sample. Cytocoarsening was run on all datasets with *P* = 10 passes, thresholds *T*^*d*^ = 26 and *T*^*q*^ = 26, and the max number of coarse nodes as 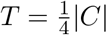, where |*C*| denotes the number of elements (coarsening candidates) of *C*.

### Accuracy and error of attributes in coarse nodes

We defined accuracy and error metrics (Fig. 2a and 2b) to evaluate the consistency of attribute values for cells assigned to a coarse node. For all of the “non super node” cells within a coarse node (e.g. those cells that were not chosen to be the representative), we predicted their attributes to be the same as that of the super node representative. The error and accuracy metrics between the true and inferred attribute labels of cells are defined as

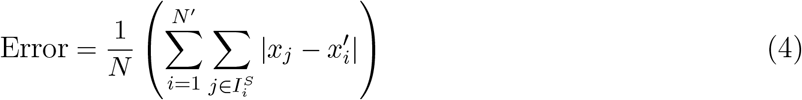

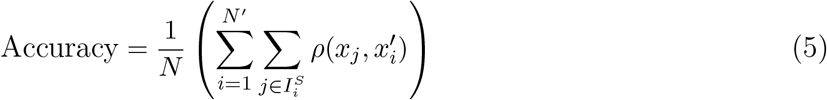

where *ρ*(*x, y*) returns 1 if *x* and *y* are equal and 0 otherwise.

**Fig. 2.**
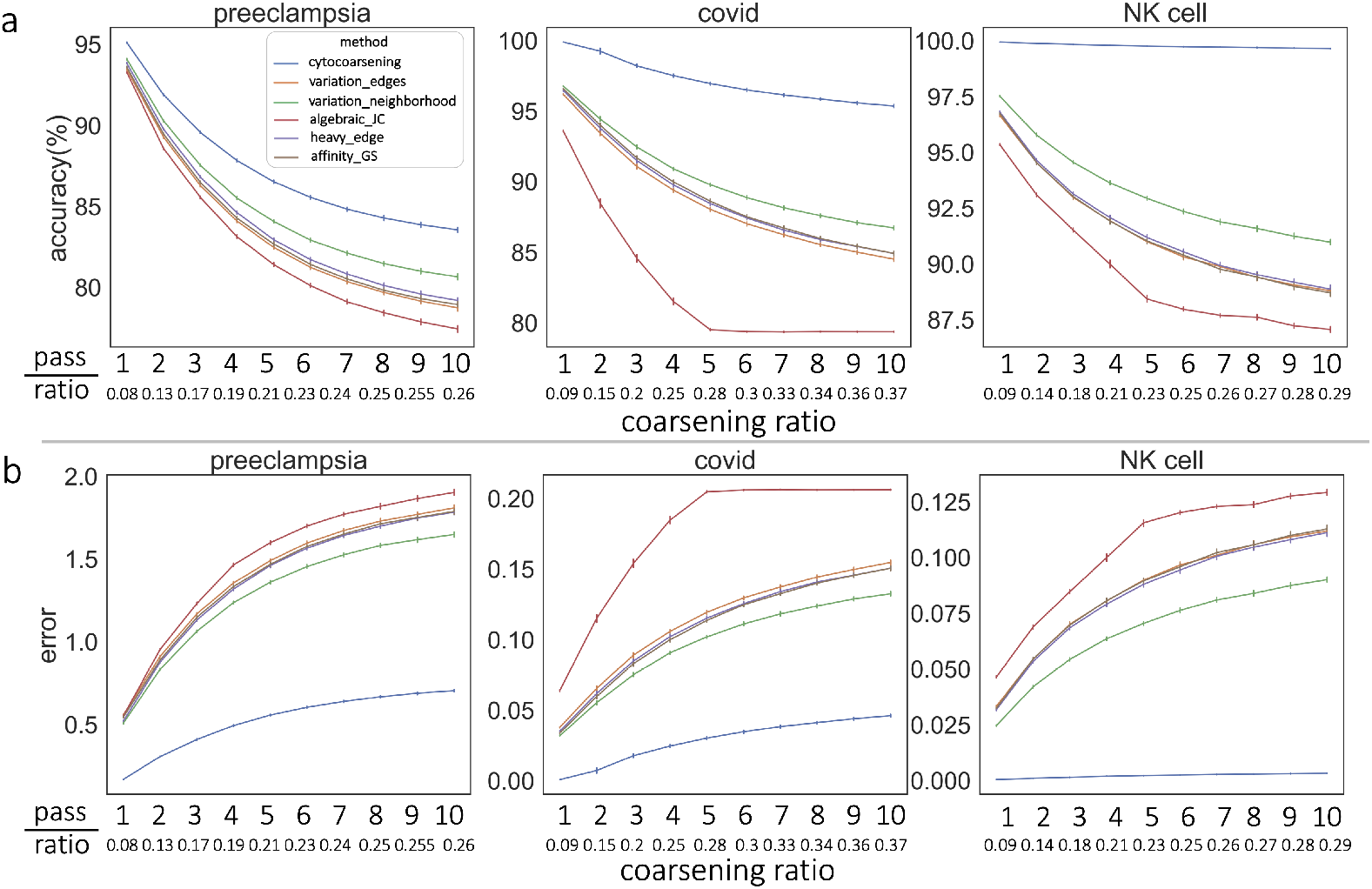
Attribute Consistency of Coarse Nodes. Accuracy (**a**) and error (**b**) metrics were used to evaluate the similarity of attributes within each coarse node. Cytocoarsening (blue) excels in accuracy and error at maintaining consistent attributes within coarse nodes across datasets.

Across datasets and coarsening ratios, Cytocoarsening exhibited superior performance, followed most closely by the variation neighborhood method. We note that the continuous attribute labels of cells in the preeclampsia dataset make the task more challenging than predicting binary attributes.

### Quantifying attribute and original feature variation across the coarse graph

Given the graph Laplacian 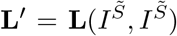 corresponding to the coarse graph 𝒢′ and the coarse attribute vector 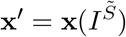, the normalized Laplacian quadratic form 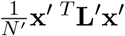 (where *N* ^′^ is the number of coarse graph nodes) summarizes the alignment between structure and attributes. Since the Laplacian quadratic form is small for vectors where neighboring nodes have similar vector entries, the quadratic form will be small if alignment is good (Fig. 3a). Similarly, we can quantify the overall variation in the features over 𝒢′ (Fig. 3b) as 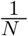 trace(**X**^′ *T*^ **L**^′^**X**^′^), where 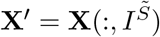 is the coarsened feature matrix.

**Fig. 3.**
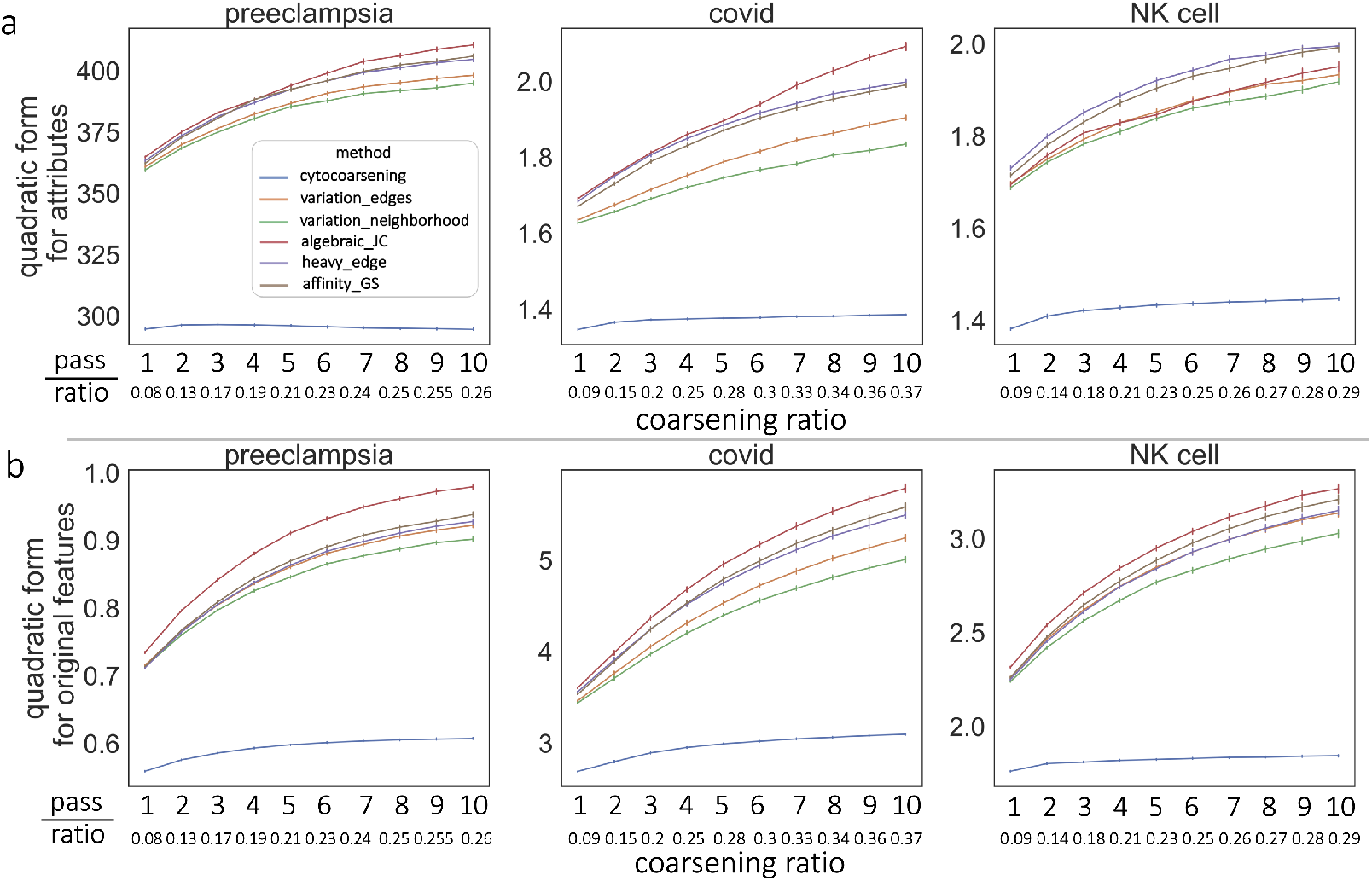
Evaluating Variation of Attributes and Original Features on 𝒢′. We used the Laplacian quadratic form on the coarse graph 𝒢′^′^ to quantify the variation of the attributes (**a**) and the original features (**b**) over 𝒢′ as a function of the extent of graph coarsening (horizontal axis). Cytocoarsening (blue) achieves by far the lowest values for both attributes (**a**) and original features (**b**) across coarsening ratios.

A good coarsening strategy would produce low values for the Laplacian quadratic forms for both attributes and in the features used to construct the original graph, implying those vary smoothly over the graph. Results across the three datasets in Fig. 3 reveals cytocoarsening produces the lowest values for both attributes (a) and original features (b) for all coarsening ratios, suggesting the cytocoarsening faithfully encodes such information.

### Coarse graphs can be used as input to MELD

To see that we would reach the same biological conclusions by analyzing G and 𝒢′, we used both of these graphs as inputs to MELD^7^ and compared the results. Given binary attribute values {0, 1}, MELD returns a list *M*, where *M*_*j*_ is the probability that node *v*_*j*_ has an attribute value of 1. We therefore binarized the returned MELD score for a node as 1 if the for node *j, M*_*j*_ *>* 0.5 and assigned it a 0 otherwise. Let **m**^*coarse*^ denote the vector of coarse graph MELD scores. We assigned all nodes within a super node *S*_*j*_ to have the same MELD score as the super node representative. Notationally, then, we have 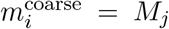 whenever node *v*_*i*_ is in the *j*^*th*^ super node. Let **m**^orig^ denote the vector of MELD scores of the original graph. We then defined two measures to quantify the similarity and correctness of the MELD results obtained for G and 𝒢′: first, *Acc*_MELD_ for accuracy. The accuracy metric quantifies the correctness of the MELD score in the coarse graph, defined as

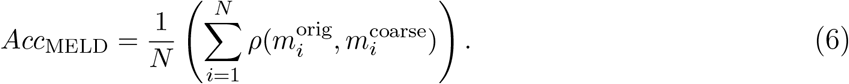

Here, *ρ*(*x, y*) returns 1 if *x* and *y* are equal and 0 otherwise. The results shown in Fig. 4b show that cytocoarsening has the highest MELD score correctness in the coarse graph when setting the smoothness parameter to the default of *β* = 1. We note that the attributes for preeclampsia dataset were dichotimized into early and late pregnancy. Although the other methods achieved accuracies above 0.9, cytocoarsening consistently achieved the highest results across datasets with both discrete and continuous attributes. Next, we computed *Corr*_MELD_, which is the Pearson correlation ^a^ between MELD scores of the coarsened graph and those of the original graph (Fig. 4a).

**Fig. 4.**
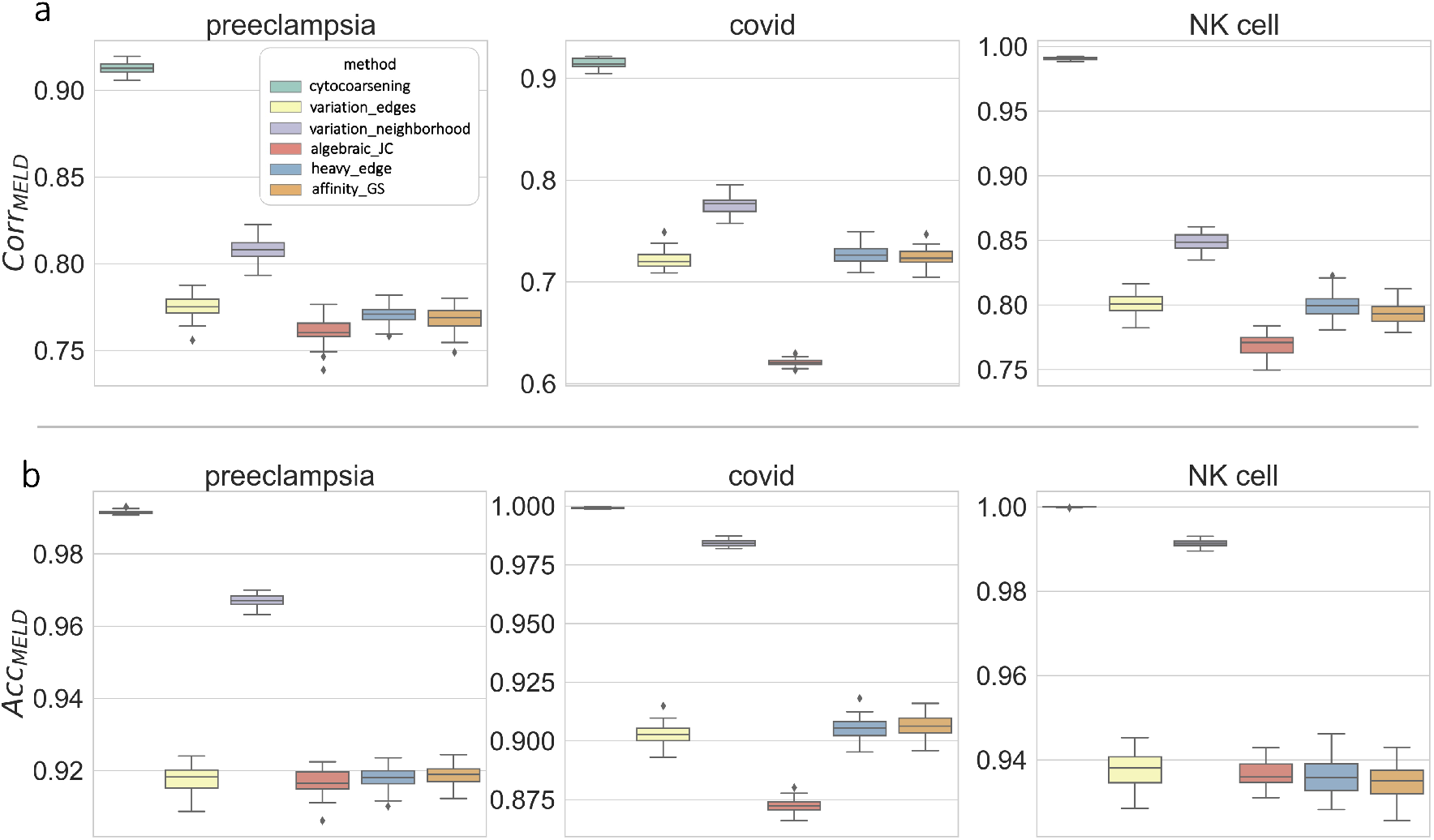
Quality of MELD Using 𝒢′ as Input. We computed metrics to evaluate the correlation (**a**) and the overall accuracy (**b**) between MELD results obtained on and 𝒢′ for six different coarsening methods and three datasets. Results suggest that cytocoarsening, followed by LV-N, produce coarse graph representations that are adequate inputs to MELD.

A high correlation implies high concordance between the MELD scores using 𝒢′ as input and those obtained using 𝒢, i.e. no critical biologically-meaningful information was lost by reducing the size of the graph. All coarsening methods achieved a reasonable *Corr*_MELD_ in all three datasets (Fig. 4a), with cytocoarsening excelling and followed most closely by LV-N.

### Sensitivity of MELD parameters in coarse graph representations

MELD has a critical parameter, *β*, which controls the smoothness or consistency of MELD scores across the graph. To study performance as a function of *β*, we varied *β* when computing MELD scores on both the original graph 𝒢 and the coarse graph 𝒢′ (we denote the parameter in each case as denoted *β* and *β*^′^, respectively). We note that due to MELD’s expensive runtime, all experiments used only 200 cells per sample. The resulting *Corr*_MELD_ scores (averaged over 30 trials) are visualized in the heatmap in Fig. 5. Cytocoarsening achieved the highest scores (denoted by stars) across datasets and combinations of *β* and *β*^′^ in 29 of the 48 comparisons (e.g. heatmap grids). The LV-N and LV-E methods are second and third in performance with a total of 12 and 11 best scores, respectively, and they perform more optimally for high values of *β* and *β*^′^.

**Fig. 5.**
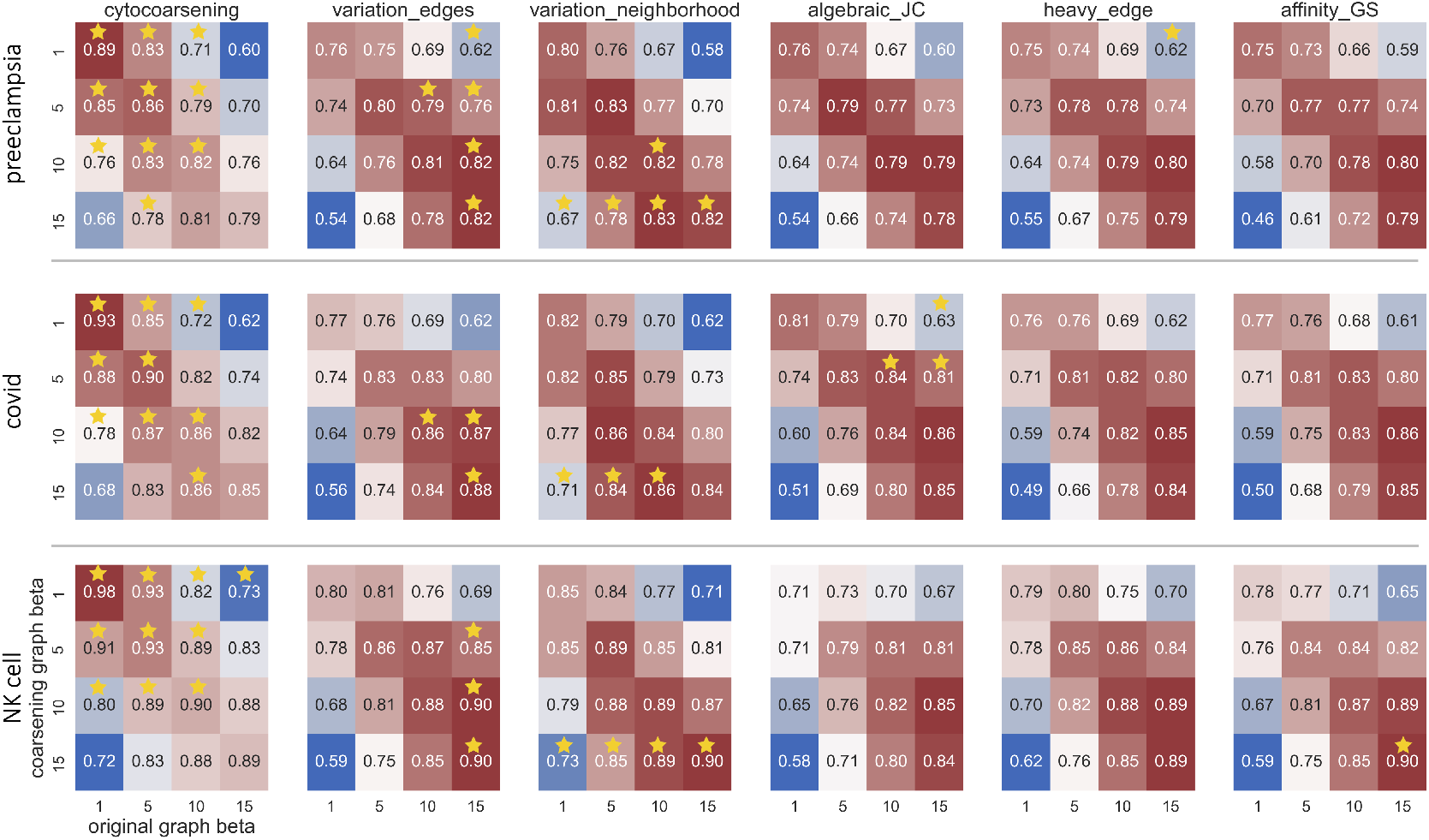
Sensitivity of MELD Results to *β* Parameter. We evaluated the effect of various combinations for values of MELD’s smoothing parameter, *β* across datasets coarsening methods. Each heatmap grid reflects the *Corr*_MELD_ obtained using 𝒢 (horizontal axis) and 𝒢′ (vertical axis) for a particular dataset, coarsening algorithm and combination of *β* parameters. A starred grid entry implies that, for that particular combination of *β, β*^′^, and dataset, the starred algorithm achieved the highest *Corr*_MELD_ score; this is frequently achieved by cytocoarsening.

### Runtime and scalability

We compared the scalability of cytocoarsening to the methods implemented to all other coarsening methods ^b^ using 1000 subselected cells from each sample. (Fig. 6). To objectively compare our multipass cytocoarsening method to existing coarsening methods, which are only one pass, we also ran cytocoarsening with a single pass. Our results show that AFF has by far the longest runtime across three datasets. Although cytocoarsening is not the fastest method, the runtime only differs slightly from the other four methods. The preeclampsia dataset is the largest in terms of patient samples and measured features and hence took the most time. In contrast, the NK cell dataset is significantly smaller and took half the time (Fig. 6).

**Fig. 6.**
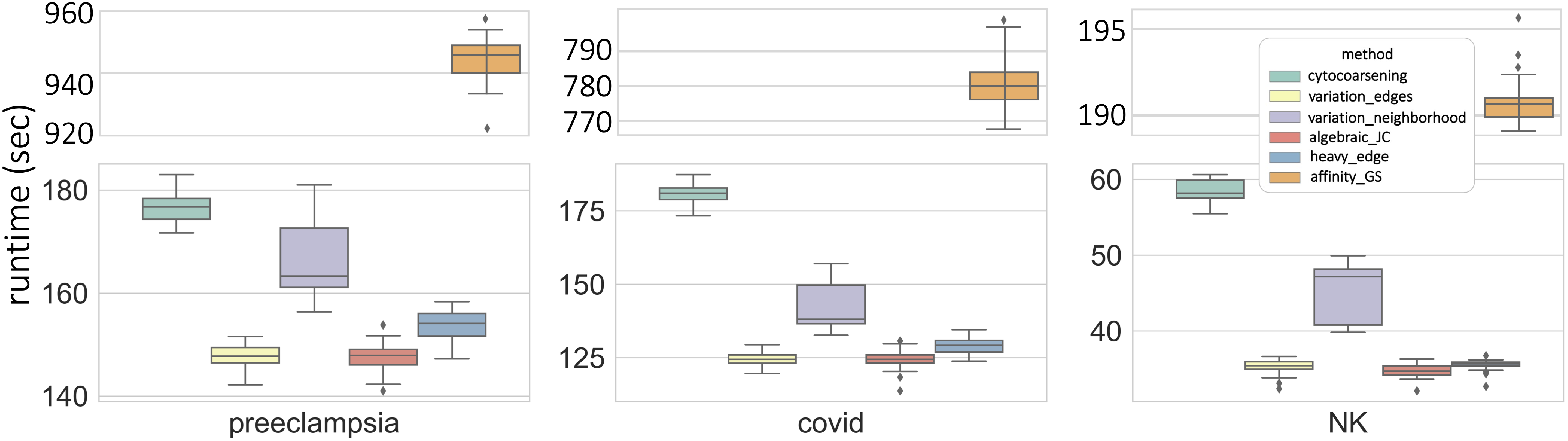
Run-Time Evaluations. Evaluating run-time of all coarsening approaches across datasets, using 1000 cells per profiled sample. Cytocoarsening has similar run-times to the other coarsening strategies, while offering increased performance in encoding attribute information.

## 4. Discussion

The cytocoarsening algorithm compresses graphs of single-cells by adapting standard graph coarsening approaches to accommodate the associated clinical or experimental cellular attributes. While existing graph coarsening approaches are optimized to create a compressed graph representation with strong *structural* similarity to the original graph, our approach uses new cost functions and a joint ranking strategy to incorporate biologically meaningful cellular information into the coarsening process. We defined several quantitative evaluation strategies to evaluate cytocoarsening and the other existing coarsening approaches on their capacity to preserve more than just structural properties of the original graph. Using three CyTOF datasets, we showed that, in comparison to other methods, the cytocoarsening method excels in grouping together cells that are both related in phenotype and in disease state or experimental condition.

Cytocoarsening is a methodological innovation towards adapting primarily structure-preserving coarsening algorithms to single-cell data with associated clinical or experimental attributes, with the aim to compress the input graph for downstream graph-based bioinformatics algorithms. However, to further increase the utility of cytocoarsening in analyzing modern multi-sample flow and mass cytometry datasets, we can modify the initial graph-construction phase for improved scalability. An area of future work is to build coarse graph representations for each sample in *parallel*, and then merge there graphs in a principled manner. Further, additional work can explore how to optimize the coarsening ratio for a particular graph. In summary, Cytocoarsening facilitates more rapid identification of phenotypically-similar cells that are likely associated with a clinical or experimental condition.

## Acknowledgments

Partial support for Emma Crawford is gratefully acknowledged from the National Science Foundation, award NSF-DMS-1929298 to the Statistical and Applied Mathematical Sciences Institute.

## Footnotes

a https://docs.scipy.org/doc/scipy-0.14.0/reference/generated/scipy.stats.pearsonr.html

b https://github.com/loukasa/graph-coarsening

